# Development and Evolution of Drosophila Chromatin Landscape in a 3D genome context

**DOI:** 10.1101/2022.11.28.518159

**Authors:** Mujahid Ali, Lubna Younas, Jing Liu, Qi Zhou

## Abstract

Chromatin states of genes and transposable elements (TEs) dictated by combinations of various histone modifications comprise key information for understanding the mechanisms of genome organization and regulation. However, little is known about the principles of their dynamic changes during development and evolution in a three-dimensional genome context. To address this, we study *Drosophila pseudoobscura*, a Drosophila model species that diverged from *D. melanogaster* about 25 million years ago. We collected 71 epigenomic datasets targeting 11 histone modification marks and 4 Hi-C datasets, and projected 15 chromatin states across four different developmental stages and two adult tissues. We estimate that before zygotic genome activation, 41% of the genome has already been deposited with histone modifications, while 20% of the rest genome switches from a ‘null’ state to an active/inactive chromatin state after the zygotic genome activation. Over two thirds of the genomic region exhibit at least one transition between different chromatin states during development. And such transitions on *cis*-regulatory regions are associated with tissue- or stage-specific formation of chromatin loops or topologically associated domain borders (TABs), as well as specific activation of gene expression. We further demonstrate that while evolutionarily young TEs are preferentially targeted by silencing histone modifications, old TEs are more frequently domesticated as TABs or specific enhancers that further contribute to the genome organization or local gene regulation. Interestingly, this trend is reversed on the newly evolved X chromosome in *D. pseudoobscura*, due to the acquisition of dosage compensation mechanism. Overall we characterize the developmental and evolutionary dynamics of Drosophila epigenomic states, and highlight the roles of certain TEs of different evolutionary ages in genome organization and regulation.

## Introduction

The highly heterogeneous sequences of the eukaryotic genome undergo dynamic epigenetic modifications that facilitate its local packaging into different states of chromatin units and global folding in the 3D nuclear space^1,2^. As a result, specialised functions can be instructed from the same genome in a spatiotemporal manner. Therefore charting the epigenomic map (e.g., patterns of histone post-translational modifications (PTMs) or DNA methylations) by high-throughput sequencing comprises a major task of consortia projects like Encyclopedia of DNA Elements (ENCODE) of human^3^ or other model organisms (modENCODE)^4^, in order to annotate the non-coding genomes and advance our understanding into the principles of genome regulation. These coordinated efforts yielded rich resources of chromatin landscapes delineated by Chromatin Immunoprecipitation Sequencing (ChIP-seq) data, targeting various histone modifications and transcription factors in highly divergent model organisms. It has now become well-established that certain combinations of PTMs show deeply conserved associations between worm, fly and human, with euchromatin (e.g., histone trimethylation at lysine 36, H3K36me3, acetylation at lysine 9, H3K9ac), constitutive (e.g., H3K9me3) and facultative (H3K27me3) heterochromatin, or cis-regulatory (H3K27ac, H3K4me1) regions (CRE)^4–7^. Nevertheless, between 34% to 68% of eukaryotic genomes, depending on the species and the numbers of analysed histone marks, were characterized with weak or no binding signals of known active or repressive histone modification marks (termed as ‘BLACK chromatin’ in one study^7^), which in Drosophila was shown to exhibit features of canonical heterochromatin (e.g., gene poor, silencing of inserted transgenes). Compared to the systematic works (e.g., Roadmap Epigenomics Project) in human and mouse, much less is known in other species about how chromatin states change across different tissue and stages throughout the development process, and it is even less clear how interspecific chromatin states evolve in response to turnovers of karyotype and repeat content^8–10^.

Besides impacting the accessibility and transcriptional status of encompassing genes, change of local chromatin states can also contribute to that of 3D chromatin architecture. This has been uncovered by the development of high throughput chromatin conformation capture (Hi-C) techniques ^11,12^. Compared to other model organisms, Drosophila have the great advantages of streamlined genomes with abundant powerful genetic tools in uncovering the debating relationship between 3D chromatin architecture vs. gene transcription^13^. Similar to other species, mitotic chromosomes of Drosophila are found to form active (A) or inactive (B) compartments, and topologically associated domains (TADs), the latter of which in Drosophila only forms upon zygotic genome activation (ZGA)^14^. TADs are hypothesised to be critical for specific and precise activation of transcription by constraining the interaction between genes and their distant enhancers^15^. However, a recent study did not find coupled large scale changes of gene expression between the highly rearranged alleles of heterozygous balancer lines of *D. melanogaster*^*13,16*^. In mouse, a series of genetic manipulations of TAD boundary (TAB) within the *HoxD* gene cluster failed to detect the pronounced expression and phenotypic changes in limbs^17^, which also questioned the causative role of TAD in shaping the gene expression.

Another advantage of *Drosophila* species is that six of their ancestral chromosome arms (termed “Muller element”) show highly conserved intrachromosomal gene content with few translocations between the elements^18^. This offers an excellent system to investigate the evolution of chromatin architecture and gene regulation in response to chromosome rearrangements. Independent fusions between the ancestral sex chromosome pair (the Muller A elements) and other autosomal elements have recurrently created sex-linked autosomes (‘neo-sex’ chromosomes), and led to turnovers of sex chromosomes ^19^. Most studied neo-sex chromosome have been found to exhibit canonical properties of ancestral sex chromosomes within a short evolutionary time, i.e., degeneration of the neo-Y^20^, and acquisition of dosage compensation mechanism on the neo-X ^21^. Here we study such a Drosophila model species, *D. pseudoobscura*, whose ancestral Y chromosome was replaced by an autosome (homologous to chr3L of *D. melanogaster*) after its homolog fused to the ancestral X chromosome (XL), giving birth to a neo-X chromosome (XR), and neo-Y chromosome^22^. While the ancestral Y chromosome shared by all other Drosophila species has fused to the dot chromosome and become an autosome ^23^ (**Figure 1a**). Here we collect 71 ChIP-seq data targeting 11 PTM marks across seven stages or tissues, among four of the tissues/stages we also collect Hi-C data. We focus on comparisons between prezygotic and ZGA stages, adult somatic and germline tissues. Besides providing a complete atlas of spatiotemporal chromatin state of genes, TEs, and CREs throughout the life cycle of *D. pseudoobscura*, we further compare it to that of *D. melanogaster*, and address how the chromatin architecture evolves in response to dramatic changes of chromosome composition.

**Figure 1.**
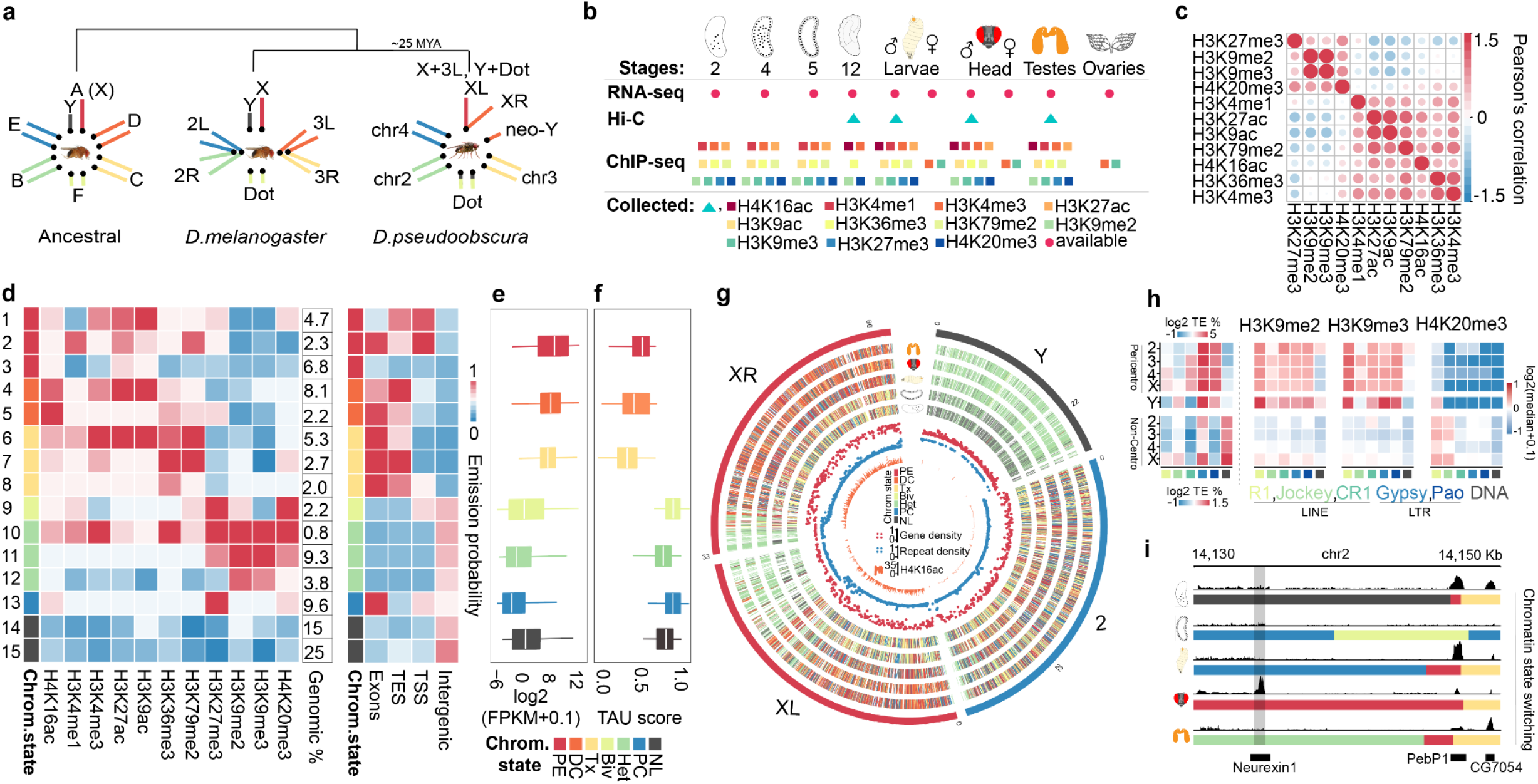
An atlas of chromatin states of *D. pseudoobscura*. **(a)** Karyotypes of Drosophila ancestor, *Drosophila melanogaster* and *D. pseudoobscura* species. About 25 million years ago, the homolog of *D*.*melanogaster* chr3L fused with that of chrX and became a neo-X (chrXR), and the homologous chr3L became a neo-Y. In addition, the ancestral chrY fused with the dot chromosome and became an autosome (Y-to-dot) in *D*.*pseudoobscura*. **(b)** The ChIP-seq and Hi-C datasets used in this study. From left to right, stage 2 (nuclear cycle 8-9), stage 4 (nuclear cycle 12), stage 5(nuclear cycle 14), 3rd instar larvae, virgin adult head (3-5 days old), virgin adult testis (3-5 days old) and virgin ovary (3-5 days old). **(c)** Pearson’s correlation between different histone post-translational modification marks. The color and circle size are scaled to the correlation coefficient, and the red color indicates a positive correlation, while the blue color indicates a negative correlation. **(d)** The 15-state chromHMM model using results of the testis as an example, other samples are shown in **Extended Data Fig. 1**. The 15 states are classified into seven biologically meaningful categories, including promoter and enhancer (PE), dosage compensation (DC), transcriptional (Tx), bivalent (Biv), heterochromatin (Het), Polycomb (PC), and Null states respectively. The numerical column represents the genomic coverage of each chromatin state. The right side heatmap shows the chromatin state enrichment within each genomic feature. **(e)-(f)** Box plots show the expression levels or the tau values of genes of different states. The higher the tau value is, the more specific the expression of the gene is. **(g)** Each gene in the genome is labeled with the associated chromatin state in chromosomes 2, XL, XR, and Y. Each track are labeled with a given stage/tissue picture. Inner red, blue, and orange tracks represent gene density, repeat density, and H4K16ac IP/IN enrichment, respectively. **(h)** The leftmost heatmap shows enrichment of major repeat types (R1, Jockey, CR1, Gypsy, Pao, DNA) in each chromosome’s pericentromeric and non-centromeric region while the chrY as a whole in *D*.*pseudoobscura*. The three right heatmaps show the normalized enrichment levels of H3K9me2, H3K9me3, and H4K20me3. **(i)** Expression pattern and associated chromatin state of the ‘*Neurexin1*’ gene across development.

## Results

### An atlas of chromatin states of *D. pseudoobscura*

To fully annotate the chromatin state, we first improve the published *D. pseudoobscura* genome assembly (UCI_Dpse_MV25) by incorporating our Hi-C data, and particularly resolve the sequences of pericentromeric regions and the partial neo-Y chromosome enriched for abundant and complex repeat sequences, with 96% of the estimated genome size now anchored into six chromosomal sequences. Only 2% of the current genome has assembly gaps, and slightly higher repeat content (∼28 vs. 22%, excluding the neo-Y) is annotated than the previously reported assemblies (**Figure 1a, Supplementary Table S1**). We then target six active histone modification marks (H3K4me1/3, H3K27ac, H3K9ac, H3K36me3, H3K79me2)^4^, and one Drosophila dosage compensation mark H4K16ac ^24^, as well four repressive marks (H3K27me3, H3K9me2/3, H4K20me3)^4^. Their normalized binding strengths along the genomic region exhibit an expected significant association (*P* < 0.05, Pearson’s correlation test) between different active marks or between repressive marks that suggest their co-binding at the same region (**Figure 1c, Extended Data Fig. 1a**). Other evidence supporting the high-quality of our data come from individual mark’s characterized distribution along the gene body (e.g., the bias toward 3’ end of active genes of H3K36me3), and distinct binding levels between active vs. inactive genes (**Extended Data Fig. 1b**). We consistently find an enrichment of active marks (e.g.,H3K4me3 specifically at transcriptional start site or TSS) on active genes, and that of repressive marks (e.g., polycomb mark H3K27me3, and the previously uncharacterized H4K20me3) on inactive genes.

To better reflect the combinatory binding patterns of these PTM marks, we demarcate the entire genome into 15 chromatin states (**Figure 1d**), which are annotated as seven state categories of Promoters and Enhancers (PE), Dosage Compensation (DC, available after embryonic stage 12), Active Transcription (Tx), Bivalent (Biv), Heterochromatin (Het), Polycomb (PC), and Null respectively, based on the reported functional associations of individual marks (**Extended Data Fig. 1c**). These state categories show biased distributions toward or at TSSs (PE), TESs (Tx state characterised with H3K36me3, H3K79me2), exons (Tx, DC and PC), and intergenic regions (Het, mostly on TEs, see below), and their encompassing genes correspondingly exhibit significant (*P* < 0.05, Wilcoxon test) differences between categories: genes of active states (PE, Tx, and DC) are transcribing at a higher level, and more likely to be housekeeping genes (i.e., less likely to be tissue-biasedly expressed) than the rest inactive state categories (**Figure 1e-f, Extended Data Fig. 1d-e**). In addition as expected, the active chromatin (A) compartments inferred by Hi-C data (on average 78% of the A compartment regions) are enriched for active state genomic regions, while inactive or B compartments (84%) are enriched for repressive state genomic regions, except that in the prezygotic stage, the Null state regions are similar between the two types of compartments (**Extended Data Fig. 1f**).

The percentage of the genome that is annotated as the Null state, or exhibits no or weak bindings of all investigated PTM marks, decreases from 61% at embryonic stage 2 (or mitotic cycle 9) to 40% in adult testis. The highest percentage of ‘Null’ genes at stage 2 is consistent with the expectation that the majority of chromatin is at a ‘naive’ state with few PTMs^25^. Nevertheless, we find that a significantly higher percentage (*P* < 0.05, Chi-square test, 38.1%) of the previously reported maternal genes relative to the genome background in *D. pseudoobscura* ^26,27^, whose transcripts are deposited into the eggs by mother, reside in a PE chromatin state genes at stage 2. While only 22% of the reported zygotic genes that only become activated after ZGA are in the same state at the same stage (**Extended Data Fig. 2a**). As expected, over 40% of the zygotic genes or maternal-to-zygotic transition (MZT) genes reside in a Tx state at the onset of ZGA (stage 5). In particular, using the normalized PTM binding levels of active/inactive genes in the adult head as a baseline (**Supplementary Methods**), we estimate 35% of the total genes, and an excess (66%, *P* < 0.05, fisher exact test) of the reported maternal genes are bound by one of the H3K4me3, H3K36me3 or H3K79me2 at their TSSs or coding regions; and only 9% of the total genes and 13% of the zygotic genes are bound by H3K27me3. This suggests that consistent with mammals, Drosophila maternal genes are enriched for potentially maternally deposited active histone modifications^28^. 24% of the TEs are bound by H3K9me2/3 (**Supplementary Table S2**) at the prezygotic stage, although this could be an underestimate because of the TE regions that cannot be uniquely mapped by the sequencing reads. These results are consistent with the previous reports in *D. melanogaster* that the PC mark H3K27me3 and HP1 protein are maternally deposited into the egg^29,30^. In contrast, no or few binding signals have been detected for the enhancer marks (H3K4me1, H3K27ac) and heterochromatin mark H4K20me3 (**Extended Data Fig. 1c**). The strong positive correlations between active PTM marks, particularly between different enhancer marks, or between repressive PTM marks only become evident after the ZGA (**Extended Data Fig. 1a**). And in testis, this is further manifested as two large correlation clusters of active or repressive marks (**Figure 1c**).

We also find variations of bindings between different constitutive heterochromatin PTMs within different genomic regions, and also between different developmental stages. At a chromosome level, approximately 69% of the pericentromeric region of XR/XL is already at a Het state at the prezygotic stage, and 18% of the region seems to have undergone reprogramming to become a Null state at the onset of ZGA (**Figure 1g**). While the majority (44%) of the neo-Y chromosome sequences, and other repeats in the non-centromeric chromosome arm regions are at a Null state before ZGA. This reflects the different properties of constitutive heterochromatin at the pericentromeric vs. chromosome arm regions. A closer examination of the repeat content between the two indicates that while the pericentromeric regions are specifically enriched for long interspersed nuclear elements (LINE) R1 and CR1, long terminal repeat (LTR) elements Gypsy and Pao, the chromosome arm regions are instead enriched for DNA transposons (**Figure 1h**). The pericentromeric region, primarily the R1 elements, have already been bound by H3K9me3 at embryonic stage 2, while other pericentromeric repeats only become bound by both H3K9me2 and H3K9me3 starting from the onset of ZGA^31^. Previous studies in Drosophila and mammals^32,33^ showed an understudied PTM H4K20me3 is associated with pericentromeric heterochromatin and retroposon silencing. Interestingly, here we find that except in the *D. pseudoobscura* head tissue, H4K20me3 becomes established on chromosome arm TEs or gene bodies of silenced genes since the onset of ZGA (**Extended Data Fig. 2b**). Overall, the genome-wide characterization of chromatin states allows us to next examine how they change during development or between tissues, and associate such changes with the specific functions of genes or CREs. One such example can be seen from the gene *Neurexin 1* (*Nrx1*) (**Figure 1j**)^34,35^, which specifically transcribes in heads of both *D. melanogaster* and *D. pseudoobscura*, and regulates synaptic architecture and contributes to the regulation of learning, memory and locomotion. This gene is encompassed in a genomic region of PE state in the head tissue, while the same region in all other tissues/stages is in an inactive PC/Het/Null state.

### Dynamic changes and correlations of chromatin states, enhancers and chromatin architecture

The temporal change of epigenetic configuration is strongly associated with the regulation of gene transcription and formation of 3D chromatin architecture, but their causal relationships remain controversial ^36,37^. Our chromatin state and Hi-C data of multiple tissues and stages provide us a great opportunity to investigate this. We first compare the chromatin state of each gene across different stages and tissues of *D. pseudoobscura*. Overall, only 32% of the total genes remain in the same category of chromatin state across the developmental course, ranging from 0.1% of the total genes remained in the Biv state to 13% of the total genes in the Null state (**Figure 2a**). Genes that are constantly in a Null state are significantly (adjusted *P* < 10^−3^, Fisher’s Exact test) enriched for Gene Ontology (GO) categories of environmental perception such as ‘sensory perception of chemical stimulus’, ‘response to bacterium’, ‘perception of taste’, consistent with their tissue/stage-biased expression (**Figure 1**). While the constant Het-state genes are enriched for GOs of nuclear or cellular functions like ‘chromosome organization’, ‘meiotic structures’ and ‘chromatin condensation’, consistent with heterochromatin’s important role in nuclear organization ^38,39^. The constant Tx- or DC-state genes are enriched for various metabolism-related GOs, and finally, constant PC- and PE-state genes are enriched for organ development or cell differentiation-related GOs (**Extended Data Fig. 3**).

**Figure 2.**
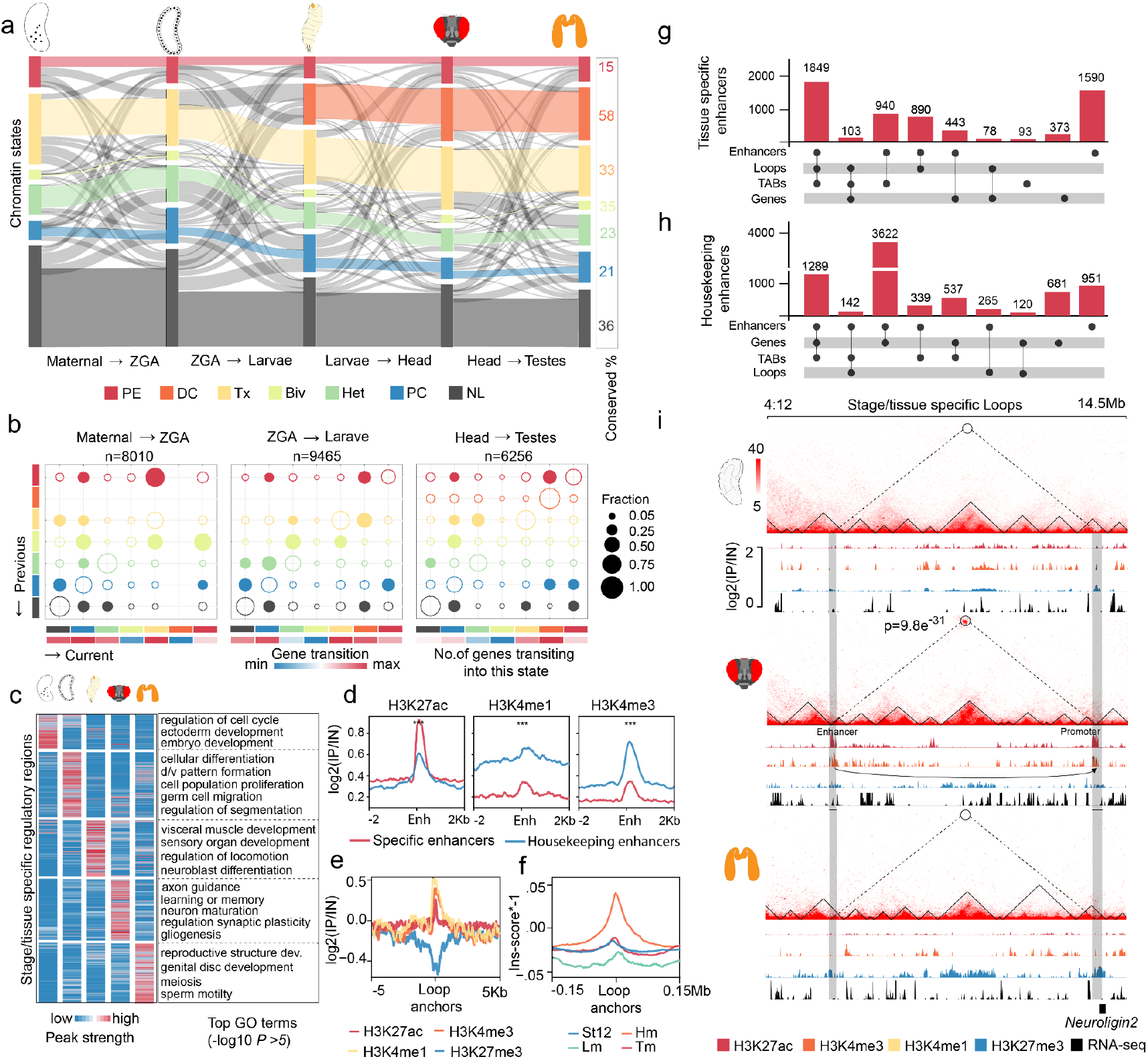
Dynamic changes and correlations of chromatin states, enhancers, and chromatin architecture. **(a)** Each color bar represents the scaled numbers of genes of each chromatin state. The colored links show genes remained in the same chromatin state across the neighboring samples. And the grey links indicate transitions of chromatin states. The numeric column on the right shows the percentage of genes of each chromatin state that remain unchanged throughout all inspected tissues/stages for the respective state.**(b)**The bubble size is scaled to the percentage of genes of certain chromatin states that show transitions from one state (x-axis) to another (y-axis) between two tissues/stages. Filled bubbles show cases when over 25% of the genes of a certain state undergo a transition, while the hollow bubbles indicate no transition or the percentage of transitions is below 25%. We show at the top of each panel also the total numbers of genes that undergo transitions, and at the bottom the heatmap indicates the numbers of genes of each type of transition. ZGA: zygotic genome activation. **(c)** Patterns of H3K27ac normalized peak binding strengths of annotated tissue-specific enhancers in a given stage or tissue. We also show the enriched GO terms of the nearby genes of the enhancers of each tissue/stage. **(d)** Metagene profiles of H3K27ac, H3K4me1, and H3K4me3 on head-specific and housekeeping enhancers. **(e)-(f)** The PTM binding patterns and insulation scores of head-specific chromatin loops anchor points. The higher the minus insulation score is, the more likely the region colocalizes with a TAB. St12: embryonic stage 12, Lm: male larvae, Hm: male head, TM: testis. **(g)-(h)** Overlap numbers of enhancers with chromatin loops, TABs, and genes for tissue-specific or housekeeping enhancers. **(i)** A head-specific enhancer with specific bindings of H3K27ac and H3K4me3, forms a specific chromatin loop with the promoter of the *Neuroligin* 2 (*Nlg2*) gene.

The rest nearly 70% of the genes undergo at least one chromatin state transition across the studied stages or tissues. In order to examine the details of transitions, we tabulate and quantify all transitions of genes between pre- and post-zygotic stages, and between adult head and testis, which respectively reflect the epigenomic transitions during ZGA and between adult somatic and germline tissues. Overall, the numbers of genes that exit the Null state or become bound by certain PTM reach the highest after the onset of ZGA (**Figure 2a**), which contributes to the largest number of state transitions occur between ZGA and larvae, together with the onset of DC (**Figure 2b**). In particular, from the prezygotic stage to the onset of ZGA, majorities of transitions are from the other states toward either the Tx or the Het/PC state. While the former is significantly (*P* < 0.05, Fisher exact test) enriched for the reported zygotic (e.g., *ftz* ^40^) or MZT genes, the latter is enriched for the reported maternal genes (e.g., *nanos* ^41^) (**Extended Data Fig. 4a**). This indicates the dynamic PTM turnovers on the maternal/zygotic genes during the ZGA. Consistently, the genes whose chromatin states are transiting toward an active state during ZGA are enriched for functional categories of ‘syncytial blastoderm’, ‘pole cell development’, ‘germ cell migration’, ‘neuroblast differentiations’ and ‘segment specification’ and so on; and those that transit toward an inactive state are enriched for ‘RNA-splicing’, ‘mitosis cycle’ and so on. (**Extended Data Fig.4b**). Between ZGA and larvae, the majorities of transitions are toward the Tx or DC state, the latter of which shows few variations between adult tissues (**Figure 2a-b**). And the genes that show a different chromatin state between head and testis reflect the specialized functions of the two adult tissues: genes that are in an active state in head but an inactive state in testis are enriched (adjusted *P* < 0.00212, Fisher’s Exact test) for GO terms of ‘sleep’, ‘short-term memory’, ‘courtship behavior’, while genes showing the opposite pattern of chromatin states are enriched for ‘spermatogenesis’, ‘mating behavior’, and ‘gamete generation’ and so on. These results together indicate the tissue- or stage-specific changes of genes’ chromatin states are highly correlated with their specific functions.

It was recently suggested that gene expression is regulated independently by the TADs preventing the spurious contacts, as well as the tethering elements facilitating chromatin loops between active enhancers and promoters^42^. To provide insights into the complex relationships between chromatin states, TADs, and interacting CREs that manifest as chromatin loops, we first annotate all the candidate enhancers based on their specific (a total of 43% of the total enhancers) or shared (57% enhancers, termed as ‘housekeeping enhancers’) binding peaks of H3K27ac across all the investigated stages/tissues (**Figure 2c**). Our accuracy of the enhancer annotation is supported by the enriched GOs of their nearby or likely the targeted genes that are highly reflective of the respective stage/tissue (**Figure 2c**). For example, genes nearby the head-specific enhancers are enriched for (adjusted *P* < 0.0035 Fisher’s Exact test) ‘axon guidance’, ‘learning or memory’, and those nearby testis-specific enhancers are enriched for ‘meiosis’ and ‘sperm motility’. In addition, genes nearby specific enhancers expectedly have a consistent specific gene expression pattern compared with those nearby housekeeping enhancers (**Extended Data Fig. 5a-b**). And specific or housekeeping enhancers are differentially enriched for the motifs reported previously for the two (e.g., dref, rpd3 motif for housekeeping enhancers, and dsx,tj motif for developmental enhancers) in *D. melanogaster* (**Extended Data Fig. 5c**). We also annotate the TABs and chromatin loop anchors across samples. 32 to 36% of the total TABs are conserved across stages/tissues, 35 to 41% of total TABs are specific to, and 25 to 29% have shifted in certain stages/tissues (**Supplementary Methods, Extended Data Fig.5d**). While chromatin loops are characterized with enrichment of enhancer PTM markers H3K27ac and H3K4me1, and promoter mark H3K4me3, but with a depletion of polycomb mark H3K27me3 (**Figure 2e**). Tissue-specific chromatin loop anchors also specifically exhibit high minus insulation scores, i.e., frequently overlap with specific TABs (**Figure 2f**). It is noteworthy that H3K4me1 is also reported to be enriched on tethering elements that facilitate long-range promoter-enhancer contacts. Thus it is possible that some tethers might also co-localize with some of the specific loop-anchors here, although they remain to be functionally characterized in future. These data together indicate that chromatin loops reflect strong specific enhancer-promoter interactions that overlap with specific TABs.

Intriguingly, we find distinct patterns between specific and housekeeping enhancers regarding their patterns of PTM bindings and co-localizations with TABs. Housekeeping enhancers show significantly (*P* < 0.05, Wilcoxon test) lower binding strengths of H3K27ac, but higher binding strengths of H3K4me1 and H3K4me3 compared to the specific enhancers across all investigated stages and tissues (**Figure 2d, Extended Data Fig. 6a**). Between 20% to 38% of the tissue-specific enhancers, depending on the tissue or stage (**Extended Data Fig. 6b**), co-localize with the chromatin loop anchors or TABs that are specific to the same tissue; while only 16% to 22% of the tissue-specific genes co-localize with the specific loop anchors and TABs (**Figure 2g**). In contrast, only 4% to 5% of housekeeping enhancers co-localize with housekeeping TABs and loop anchors, and majority (52%) of these enhancers only co-localize with housekeeping genes (**Figure 2h**). This is further supported by the pattern that stage/tissue specific TABs have significantly higher normalized binding strengths than those of housekeeping TABs, consistent with features of specific enhancers (**Extended Data Fig. 6c, Figure 2d**). These results together indicate that interactions between spatiotemporal specific enhancers and promoters, but not housekeeping enhancers, may contribute to the specific chromatin architectures. An example is shown in **Figure 2i**, that a head-specific enhancer that exhibits specific bindings of H3K27ac and H3K4me3, forms a specific chromatin loop with the promoter of the *Neuroligin* 2 (*Nlg2*) gene specifically transcribing in the head. *Nlg2* interacts with *Nrx1* and participates in synapse formation and growth, as well as regulation of learning and memory ^43^.

### Transposable elements play both regulatory and structural roles in shaping the chromatin architecture

Among the annotated enhancers, 10% are overlapped with TEs, suggesting that they have likely been co-opted to regulate specific gene expression accompanied by their changes of chromatin states (**Figure 1g, 2a**). Before further characterizing the potentially functional role of TEs, we first chart their changing transcriptomic and epigenomic landscapes to gain a genome-wide view. All TE families in total comprise 39% of sequences of the current genome assembly of *D. pseudoobscura* (**Supplementary Table S2**). Majorities, but not all of the LINE (on average 21% among stages/tissues), LTR (36%) elements are in a Het, PC, or a Null state, while only 13% of the DNA transposons are one of the three repressive states (**Figure 3a, Extended Data Fig.7a-c**). Specifically, 20% and 7% of all TE copies (**Figure 3a**) respectively reside in the Null and Het state throughout all the stages/tissues, and majority of them are located in pericentromeric regions and form the constitutive heterochromatin. These TEs also exhibit strong interactions between pericentromeric regions of different chromosomes (**Extended Data Fig. 7d**), indicating stable clustering of centromeres of mitotic chromosomes of Drosophila species across different mitotic cell types^44^.

**Figure 3.**
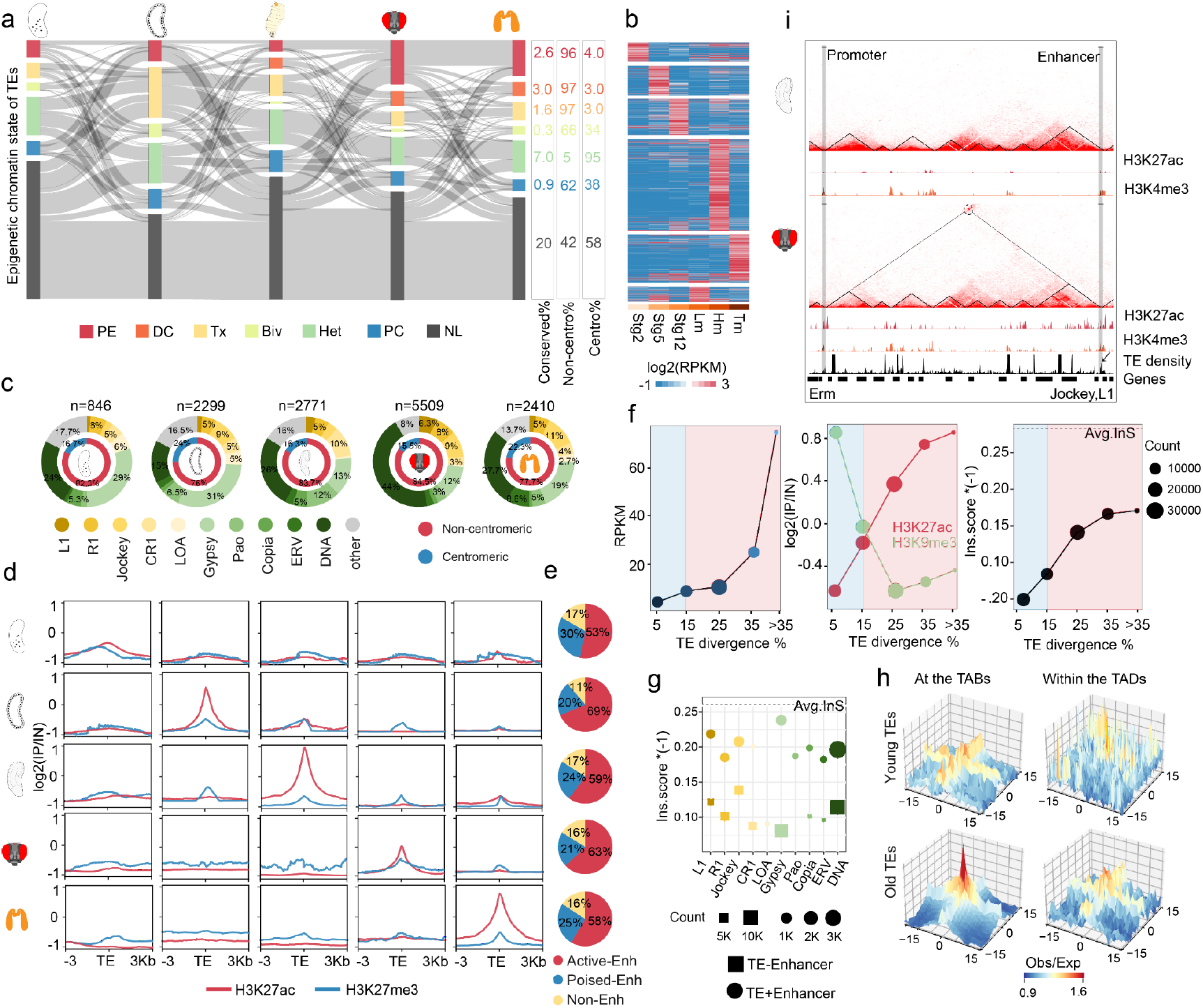
Transposable elements play both regulatory and structural roles in shaping chromatin architecture. **(a)** Chromatin state transitions of TEs across development. The numbers on the right respectively show the percentage of TEs that maintain their chromatin state across stages/tissues, and the percentages of pericentromeric or chromosome-arm TEs. **(b)** TE expression across development. Each cluster shows the tissue-specific expression patterns, and detailed expression of TEs subtypes is present in **Extended Data figure 8c. (c)** Composition of stage/tissue-specific TEs from figure b. **(d)** Enrichment of enhancer mark H3K27ac and polycomb mark H3K27me3 on stage/tissue specifically expressed TEs identified in figure b. **(e)** Composition of active, poised, or non-enhancers on stage/tissue-specific TEs **(f)** The associations of expression levels of TEs, H3K27ac/H3K9me3 binding strengths, minus insulation score values in head tissue with the divergence levels of TEs from the consensus sequences, or the age of TEs. The higher minus insulation shows a stronger TAD border. **(g)** Minus insulation scores of head-specific old TEs: circles are the TEs overlapped with enhancers, while square boxes represent the other TEs. **(h)** Left column: Long-range interactions (from 300Kb to 5Mb distance range) of young vs. old TEs present at the TAD borders, right column: same as previous but for the TEs present within the TADs. **(i)** A head-specific enhancer-like TEs (Jockey, L1) bindings of H3K27ac and H3K4me3, forms a specific chromatin loop with the promoter of *the ERM* gene.

On the other hand, substantial numbers of and a comparable percentage (65%) of TE copies relative to that of genes, undergo at least one transition of chromatin state throughout development; and only a few of them are attributed to transitions from/to the PC state. The major transitions occur between the Null vs. other states, particularly during ZGA. 5% of all TE copies exit the Null state and enter the Tx state at the onset of ZGA and reverse back to the Null state after ZGA, that is, they are specifically transcribing at ZGA (**Figure 3a, Extended Data Fig. 8a**,**b**). The other prominent transition involves those from the Null and other states into the PE state specifically in the head, which has the largest number of PE-state TEs among all tissues/stages (**Figure 3a**). These specific active states of TEs are further reflected on their spatiotemporal transcription patterns (**Figure 3b**). Between 846 to 5509 TE copies that are specifically transcribed in specific tissues or stages, and majorities of them are located in the non-centromeric chromosome arm regions (**Figure 3c**). In particularly, we identify large numbers of Gypsy elements, and several subfamilies of DNA transposons (e.g., Maverick, hAT) that specifically transcribe at ZGA; and large numbers of L1, R1 LINE elements, and CMC-EnSpm DNA transposons that specifically transcribe in the head (**Figure 3b, Extended Data Fig. 8c**). 53% to 69% of these specifically transcribing TEs are bound by H3K27ac, and between 20% to 30% of them are bound by H3K27ac and H3K37me3 specifically in the same tissue, which likely act as specific active enhancers or poised enhancers (**Figure 3d-e**). This is further supported by the consistent transcription pattern (**Extended Data Fig. 9a**) and the enrichment of relevant GOs of genes nearby these TEs. For example, genes nearby TE specifically expressed in heads are enriched for functional categories of “learning or memory”, “CNS development”; while those nearby TEs expressed in testes are enriched for GOs of “sperm motility” and “meiosis”(**Extended Data Fig. 9b**). These results suggest some TEs that exhibit spatiotemporal changes of chromatin state and transcription have likely been domesticated to regulate specific gene expression as enhancers. This gradual process of ‘TE domestication’ is also reflected by the strong correlation between their evolutionary ages vs. their transcription levels, and normalized binding levels of PTM marks. In general, the younger (measured by their sequence divergence levels from the consensus sequences) the TE copies, the less likely (*P* < 0.05, Pearson’s correlation test) they are transcribing, or bound by the enhancer mark H3K27ac, but more likely to be silenced by H3K9me3 or to be a poised enhancer bound by H3K27me3 (**Figure 3f, Extended Data Fig.9c**). These results indicate that young TEs are initially well controlled for their transposition activities by heterochromatic PTMs, and during evolution, some of them diverge in their sequences and acquire regulatory functions and exhibit bindings of active enhancer PTMs.

Besides such regulatory functions, TEs can also play an important role in shaping the chromatin architecture, as suggested by previous studies in mammals^45–49^. We find in *D. pseudoobscura* that old and domesticated TEs are more likely than the younger ones to coincide with the TABs, indicated by the formers’ similar levels of insulation scores with those of genome-wide TADs (**Figure 3f**). And the enhancer-like TEs (**Figure 3d**), particularly the Gypsy elements exhibit a significantly (*P* < 0.05, Wilcoxon test) higher minus insulation scores, or much more likely than the non-enhancer TEs to be coinciding with TABs (**Figure 3g**). Old TEs also exhibit specific interactions identified by the Hi-C data, while young TEs or TEs that reside within the TADs do not show such interactions (**Figure 3h**). In fact, 9% to 13% tissue specific chromatin loops overlap with the active enhancers derived from TEs (**Extended Data Fig. 9d**). One example that shows how the TEs exert their regulatory and structural functions is shown in **Figure 3i**. A chermic TE copy of Jockey and L1 acts as an enhancer, and forms a tissue specific chromatin loop with the promoter of the gene *erm* and likely specifically activates its expression in the head. *Erm* was reported to be contributing to the development of neural stem cells of the larvae brain in *D. melanogaster* ^50^.

### Evolution of chromatin state and regulatory elements between *D. melanogaster* and *D. pseudoobscura*

We finally set out to address the evolution of chromatin state of genes and regulatory elements between the two Drosophila species, particularly in response to their sex chromosome turnovers (**Figure 1a**). On average among different tissues and stages, 57% of the orthologous genes on the homologous autosomes of both species reside in the same chromatin state across the investigated tissues and stages, while this number decreases to 42% when comparing the chrXR of *D. pseudoobscura* to the homologous chr3L of *D. melanogaster* because of acquisition of DC mechanism on chrXL (i.e., transitions of other states into DC state) (**Figure 4a**). Of each state, Tx genes exhibit the highest level of interspecific conservation, while Biv genes seem to have undergone the most dramatic interspecific changes. And genes that undergo interspecific transitions of chromatin states are more likely to be tissue-specifically transcribed genes (**Extended Data Fig. 10a**). The major interspecific differences of chromatin state of genes on autosomes are derived from the *D. pseudoobscura* genes in Null or PC state with an *D. melanogaster* ortholog in another state, while those between chrXR and chr3L are from evolution of DC (**Figure 4a**). Genes that transit into a DC state on chrXR are enriched for *D. melanogaster* orthologous genes of active (PE and Tx), as well as Null state genes.

**Figure 4.**
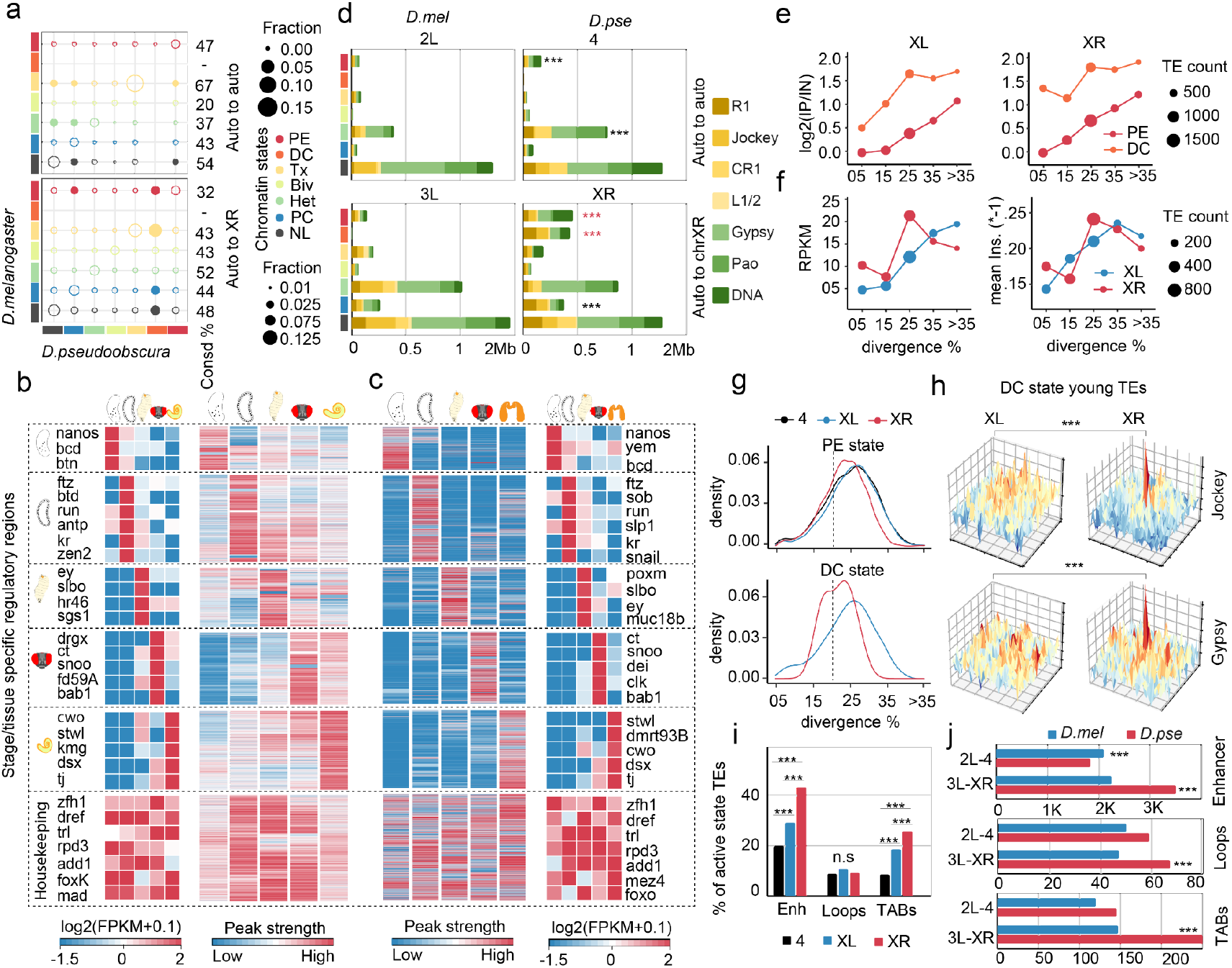
Evolution of chromatin state and regulatory elements between *D. melanogaster* and *D. pseudoobscura*. **(a)** Diversity of chromatin states in head-tissues between orthologous genes of *D. melanogaster* (x-axis) and *D. pseudoobscura* (y-axis), the color bar represents chromatin state. The upper panel shows comparisons between species on the autosomes, and the lower panel shows the comparison of *D. melanogaster* chr3L vs. *D. pseudoobscura* chrXR, numbers on the right show the percentage of genes of each state that remain conserved between species. The scaled filled circle shows the cases of chromatin state transitions if the total involved genes are over 2% of the total genes in the current state of that chromosome. Hollow circles indicate the conserved genes or transitions involve below 2% of the total genes. **(b)** Each (C1-6) cluster from left to right shows the tissue specific expression patterns of TFs predicted to bind the tissue specific enhancers of the certain stage or tissue in *D. melanogaster* or in *D. pseudoobscura* (**c**). The right panel shows the binding patterns of H3K27ac peaks that are used to annotate the tissue/stage-specific enhancers. **(d)** Comparison of composition of TEs of each chromatin state in the head between the homologous autosomes (the upper panel) and chr3L vs. chrXR between the two species (the lower panel). **(e)** chrXR but not chrXL are enriched for young DC-state associated TEs. x-axis shows the divergence% of TE, y-axis shows the enrichment level of certain age of TEs. **(f)** Left panel; expression levels of DC-associated TEs in chrXL and chrXR represented in blue and red, respectively. The x-axis is TEs divergence levels (%) from the consensus sequences, and the y-axis is the mean expression level in the head. Right panel: the association of minus insulation scores of the TEs with their divergence levels from the consensus sequences. **(g)** Age distribution of PE- and DC-associated TEs in the head along chr4, chrXL, and chrXR in black, blue and red, respectively. **(h)** Long-range interactions (from 300Kb to 5Mb distance range) of DC-associated young Jockey (upper panel) and Gypsy (lower panel) elements in chrXL (left column) and chrXR (right column), (P-value < 0.05 Wilcoxon test) in the head. **(i)** Percentage of active state TEs overlapped with head-specific enhancers, loops, and TABs(p-value <0.05 Wilcoxon test) on chr4, chrXL and chrXR represented with black, blue and red color respectively. **(j)** Colocalization of active state TEs from *D. melanogaster* (in blue) to *D. pseudoobscura* (in red), autosome to autosome (2L to 4) vs. autosome to sex chromosome (3L to chrXR) to the head-specific enhancers (upper panel), loops (middle panel) and TABs (bottom).

For enhancers, although overall 82% of the *D. melanogaster* or 76% of the D. pseudoobscura candidate enhancers (compared to 76% of the *D. melanogaster* genes) have orthologous sequences in the other species, this number decreases to only 40% and 33%, if we condition on the enhancers share the same pattern of tissue/stage specificity. This suggests that the turnovers of enhancers between species are more often attributed to those of spatiotemporal epigenetic changes than those of genomic sequences. Across the investigated tissues/stages, housekeeping enhancers (**Extended Data Fig.10b**) exhibit the highest level of conservation, with 92% of the *D. melanogaster* housekeeping enhancers having their orthologous sequences in *D. pseudoobscura* also as housekeeping enhancers. By contrast, this number decreases to only 65% for the enhancers specific to the early embryonic stages (for other tissue/stage, this percentage ranges between 72% to 81%, **Extended Data Fig. 10c**). This is consistent with the expected stronger evolutionary constraints on enhancers functioning in multiple tissues and stages, as well as the dynamic evolution of maternal transcripts reported in Drosophila^51^.

The turnovers of specific enhancers and their predicted encompassed binding motifs are strongly associated with those of their predicted binding transcriptional factors (TFs) (**Figure 4b-c**). Between 44% to 89% of the *D. melanogaster* TFs predicted to bind to specific enhancers of a certain stage or tissue are shared with those of *D. pseudoobscura*. These highly conserved TFs include *nanos* and *bcd* in the early embryonic stage 2, *ftz* and Kruppel (kr) at the onset of zygotic activation, *tj* and *stwl* in testis, which all have been reported to specifically function and transcribe in the respective tissue or stage ^52–55^. Nevertheless, a large number of TFs exhibit species-specific expression patterns (**Extended Data Fig. 10d**, **Supplementary Table S3**) and their corresponding predicted binding enhancers, with the highest interspecific diversity of TFs in testis, and the highest interspecific conservation for housekeeping TFs and in the prezygotic embryos (**Extended Data Fig. 10e**).

Interestingly, we find on chrXR of *D. pseudoobscura* an significant (*P* < 0.05, Wilcoxon test) excess of not only genes but also TEs that have turned into a DC or PE state, relative to the homologous autosomes of *D. melanogaster* or other autosomes of *D. pseudoobscura* (**Figure 4d**). This could be due to the byproduct of the spreading of the DC complex and its consequential PTM H4K16ac along the chrXR^56^. Alternatively, as shown before^21^, some newly propagated TEs on the chrXR can mediate the spread of DC. In contrast to the patterns of autosomes and the old X chromosome chrXL (**Figure 3g**), the extremely young TEs (whose divergence level with the consensus sequence is below 5%) on the *D. pseudoobscura* chrXR instead are more likely to be actively transcribing, reside in a DC state, and also more likely to be coincide with TABs (**Figure 4e-f**). Such young TEs include the previously reported Helitron elements ^21^, but also those newly identified in this work, Gypsy, Jockey and some DNA transposons, that exhibit strong specific interactions between the same type of TEs. In addition, such interactions between young TEs are absent on other autosomes of *D. pseudoobscura* or the homologous chr3L of *D. melanogaster* (**Extended Data Fig. 10f**). To further discriminate the driving forces underlying the accumulation of young and interacting TEs on the neo-X chrXR, we further compare the density of reported 21-bp binding motifs (MSL recognition element, MRE^57^) of DC protein complex, between young vs. old TEs on chrXR and chrXL. Interestingly we find that the young TEs harbor significantly (*P* < 0.05, wilcosxon test) more MRE elements than old TEs, and this pattern is specific to TEs located within the DC state and on the chrXR, but not on chrXL or TEs that are not in the DC state (**Extended Data Fig. 10g**). Given that these TEs very likely accumulate on chrXR after it acquired the DC mechanism, this reflects the ongoing evolution, rather than the initial acquisition of DC on the chrXR. That is, these young TEs might have facilitated rather than initiated the spreading of DC along the chrXR, after they have become activated by the spreading of DC. This is further supported by the pattern that the TEs resided in the DC or PE state (**Figure 4g**), but not other states (**Extended Data Fig. 10h**), tend to be significantly (*P* < 0.05, Wilcoxon test) younger than those on chrXL and autosomes. In addition, young TEs including Jockey and Gypsy tend to have specific interactions with other young elements of the same family. And this pattern is only observed on chrXR, but not on chrXL. Taken together, our results suggest some young TEs have participated in the spreading of DC on the neo-X chromosome.

## Discussions

Here we present a catalog of chromatin states of genes, TEs, and CREs across development of *D. pseudoobscura*, and the first comprehensive comparison of epigenomes between Drosophila species. We find dramatic turnovers of chromatin states on genes during the ZGA process. Before ZGA, over 60% of the entire genome is in a naive state without any PTMs, and the ZGA process involves these Null state genes transiting into a state with constitutive (Het) or facultative (Pc) heterochromatin marks. In contrast to one of few previous works that profiled the epigenomes of pre-zygotic embryos of Drosophila^25^, which might be due to the different methods of ChIP-seq experiments (**Supplementary Methods, Supplementary Table S3**), we detect over 2200 genes (**Figure 2a, Extended Data Fig. 2**) reside in a PE status as early as prezygotic embryonic stage 2. These genes are enriched at the TSS for the canonical promoter mark H3K4me3, but not for enhancer marks H3K27ac, and are significantly enriched in the reported maternal genes in *D. pseudoobscura*. It is unclear what is the function of such a pattern of poised promoter marks at the maternal genes. 49% of such genes are also MZT or zygotic genes, so it might be associated with their further activation during the later development, similar to the case in zebrafish^58^. Another interesting pattern before and after ZGA is the differential process of heterochromatinization involving different PTMs, between the pericentromeric and chromosome arm regions. Before ZGA, few chromosome arm regions except for the Y chromosomes are bound by the canonical constitutive heterochromatin marks H3K9me2/3, with variations of bindings between different families of TEs. And after ZGA, a not well studied PTM mark H4K20me3 specifically binds to the chromosome arm regions, but shows a strong tissue/stage specificity.

The dynamics of chromatin state changes on individual genes or TEs is more complex and here we specifically focus on those specific to one tissue or stage. We document large numbers of TEs, particularly those that are evolutionarily old and likely have been domesticated for various structural or regulatory functions. Such TEs include, for example, the LTR Gypsy elements enriched in the pericentromeric regions, and various DNA transposons enriched in the chromosome arm regions. On autosomes, they exhibit activation, bindings of enhancer marks H3K27ac i, and co-localization with TABs in a tissue or stage specific manner. Therefore, they are likely to drive specific expression of nearby genes, through forming a specific chromatin loop (e.g., **Figure 3i**). Given the abundance, genomic position and epigenomic status of TEs can rapidly evolve between species, these enhancer-like TEs likely significantly contribute to the diverged gene expression and chromatin architecture between the two Drosophila species.

While on the other hand, TEs on the recently evolved sex chromosome exhibit an opposite pattern regarding their evolutionary age in contrast to those on autosomes. As a consequence of spreading of DC, many TEs become reside in a hyperactive chromatin environment mediated by H4K16ac, and possibly gain the opportunity to form interactions with each other during the DC process. This pattern is only observed on the younger chrXR but not on chrXL. And such young-interacting TEs are specifically enriched for the binding motifs for the DC protein complex, suggesting another way of TEs contributing to evolution of chromatin architecture is through facilitating the spreading of DC complex throughout the entire chromosome by bringing distant genomic regions into close contact.

## Experimental Methods

### Genome assembly and Y-linked sequence annotation

The new *Dpse* assembly was built from the combination of a female assembly (UCI_Dpse_MV25) and Y-linked contigs from a male assembly (UCBerk_Dpse_1.0). To obtain an intact Y chromosome, Hi-C library reads were used as input data for 3D-DNA (Dudchenko et al. 2017), a software pipeline designed specifically for using proximity ligation data to scaffold the Y-linked contigs. The JuicerBox (v1.9.8) was then used for visual curation. For repeat annotation, we first used RepeatModeler (open-1.0.10) to construct the consensus repeat sequence library of the Y chromosome. Then, the *de novo* library and the repeat consensus library in Repbase (Bao et al. 2015) were merged to annotate all repetitive elements using RepeatMasker (open-4.0.9). We integrated evidence of protein homology, transcriptome, and *de novo* prediction to annotate the protein-coding genes with the MAKER v2.31.10^59^ pipeline to obtain Y-linked gene models.

### Fly stocks

*Drosophila melanogaster* fly stocks were maintained at room temperature with natural light/dark cycle. For *Drosophila pseudoobscura* and *Drosophila miranda* fly stocks, we used 18°C incubators with 12 hours of light/dark cycle. We raised the flies on the Institute of Molecular Pathology (IMP) standard fly food with yeast in plastic bottles and vials.

### Embryo collection

First, we validated the staging of the embryos for given biological collection time points using small (5.5 3 7.5 cm) cages. After validating the embryo age at our collection points, we move the flies to the fly population cages three days before the embryo collection. Also, we prepared agar plates (90cm) by adding 100ml of apple and grape juice to two liters of agar medium. On the day of collection, we spread the semisolid yeast paste petri-dishes, placed them in the cages, allowed the flies to lay eggs for 20 minutes, and discarded the first four batches before collection to avoid the old embryos. After discarding the four pre-lays, we kept the 5th batch of the embryos and incubated them at an 18°C incubator for a given stage. After incubation, wash the embryos from the apple-yeast-agar plates with low-stream tap water and a paintbrush into the collection sieve. Transfer embryos to a beaker filled with a freshly prepared dechlorinating solution (3 % Sodium hypochlorite) for 3 minutes at room temperature. Wash the embryo with tap water and let them dry on tissue paper in the sieve. Then transfer the embryo to a 15mL falcon tube containing cross-linking solution (1 mM EDTA, 0.5 mM EGTA, 100 mM NaCl, 50 mM HEPES, pH 8.0). Place the tube on the shaker at 300rpm for 10 minutes at room temperature. Briefly, centrifuge the tube to pelleted down the embryo and remove the supernatant. Stop the cross-linking by adding 125mM glycine with 0.1% triton-X-100 solution and vigorously shake the tube for five minutes, and pellet down the embryo by a short centrifugation at low speed. Carefully remove the supernatant, wash the embryo pellet twice with PBS solution, remove the PBS completely, and flash freeze the tube in liquid nitrogen. We repeated this process for each batch of embryonic stages 2, 4, 5, 9, and 12 embryos until sufficiently collected embryos for input material.

### Adult tissue collection

For testes, ovary, and head samples, we sorted the virgin flies under the microscope and raised them on standard food for 3-5 days. For ovary and testes, dissected 3-5 days old virgin flies and collected 200-400 pairs of testes and ovary (samples) in cold testes extraction buffer (TEB) (10mM HEPES, 100mM NaCl,1xPBS, 1x protease inhibitors, 1mM PMSF). Wash the sample in 1X cold PBS+PI twice, fix it in 1% formaldehyde at room temperature (RT) and quench the reaction by adding 125 mM glycine for 5 mins, spin, and discard the supernatant. Next, wash the tissue pellet (from the last step) twice with a homogenization buffer (140mM NaCl, 1mM EDTA, 10mM HEPES, 0.1% triton-X100, 1x protease inhibitors, 1mM PMSF) and homogenize it in 1 mL Dounce homogenizer with a tight-fitting pestle on ice, transfer the cell suspension to the 1.5mL tube and spin it. Wash the pellet once, spin it and resuspend it in a hypotonic buffer (10% glycerol, 10 mM HEPES, pH 8, 1.5 mM MgCl2, 10mM Tris HCl, pH 7.5, 0.3 M Sucrose 0.2% Triton X-100) and keep it on the rotator for 10 min at 4°C. 5. Spin the sample, and resuspend the pellet in the lysis buffer (if observing the tissue piece, homogenize the pellet in the lysis buffer by small homogenizer) and keep it on ice for 30 mins and mix it 2-3 times. Sonicate the chromatin using ultra-ultrasonicator (peak power:175, duty factor:20, cycle/burst: 200) for 2 minutes. Spin the sample at max speed for 7 min, transfer the supernatant to the new tube and adjust the final volume with an IP buffer. For head collection, 3-5 days old flash frozen virgin flies were added to the 50mL tube containing liquid nitrogen and glass beads. Vigorously shake the tube and pour the parts of flies along the beads on the specially designed apparatus comprising four levels of different sizes of sieves course one on the top and a fine one at the bottom. The top sieve retains the large body parts and beads, two middle sieve collects the heads, and the bottom sieve contains legs and wings. Check the heads under the microscope to remove, clean, and remove if there are other body parts. Afterward, heads were collected in a 1.5mL tube and flash-frozen in liquid nitrogen. Add a small amount of liquid nitrogen to the ceramic motar, and pour the heads collected at the previous step. Grind them with the pestle to fine powder, and cover the motor top with aluminum foil if heads splash out while pressing with the pestle. Transfer the powdered sample to 1mL pre-chilled glass Dounce homogenizer. After homogenization, we used a standard method for all embryonic stages and adult tissue. For the larval sample collection, we sorted and collected the third instar larvae on ice under the microscope for each sample and species. Then, flash frozen the samples in the liquid nitrogen and stored at -80°C or proceed to homogenization lysate. Keep frozen larvae in dry ice until ready for use. Then grind the frozen male larvae with the pestle to fine powder, as we did in the head sample. Transfer the powdered sample to 1mL pre-chilled glass Dounce homogenizer

### Chromatin preparation

Thaw and wash the embryo in cold 0.3% PBT containing 1mMPMSF solution and transfer them to 1mL pre-cold glass Dounce homogenizer containing 500 uL homogenization solution. Homogenize the embryo by applying fifty strokes of a loose-fitting pestle on the ice box. Transfer the homogenized lysate to a 1.5mL tube, and remove the vitelline membrane and cell debris by brief spinning. The supernatant was removed, cold cell lysis buffed used to resuspend the pellet, and again homogenize the cells in 1mL Dounce homogenizer with the tight-fitting pestle by applying fifty strokes. Transfer the sample lysate to a low-bind 1.5mL tube, centrifuge it at 4 °C for 5 minutes, and pellet down the nuclei. Remove the supernatant, and resuspend the nuclei in cold nuclear lysis buffer (1x Protease inhibitor (halt), 1mM PMSF, 10mM tris-HCl, 1mM EDTA, NP-40 0.5%, SDS 0.1 %, 0.5% N-lauroylsarcosine), and incubate on ice for 30 minutes. Sonicate the chromatin by using an S220 Covaris ultra-sonicator with the setting of peak incident power 75W, duty factor 25%, and cycle per burst 1000 for 5 minutes in specially designed microTUBE-50s. Spin the sample at max speed for 7 min, transfer the supernatant containing the chromatin to the new tube and adjust the final volume with an IP buffer (15 mM Tris-HCl pH 8.0, 2 mM EDTA, 150 mM NaCl, 0.1% Triton X-100, 1x Protease inhibitor, 1mM PMSF). Pre-clear the chromatin for each sample, take 30uL well-mixed Dynabeads in their own tube, and place them on a magnetic rack for 1 min. Carefully remove supernatant. Add 200uL Dynabeads wash solution to each tube and vortex. Place on a magnetic rack for 1 min and carefully remove the supernatant. Repeat this step twice. Allow beads to air dry for an additional 5 minutes. To dry beads, add 500uL chromatin and rotate for at least 1-2 hours at 4°C. Place samples in a magnetic rack for 1 min on an ice box, and carefully remove and retain supernatant. Immunoprecipitation: Add 2-3uL antibody to pre-cleared chromatin and rotate at 4oC overnight. For each sample, prepare Dynabeads by aliquoting 40uL well-mixed Dynabeads to a 1.5mL tube. Wash the beads as twice and let them dry for five minutes. To dry beads, add chromatin bound to the antibody (IP-antibody complex), and rotate at 4°C for at least 2-3 hours. Place samples in a magnetic rack on ice and carefully remove the supernatant. Wash beads three times with 1mL normal RIPA buffer and two times with high salt RIPA buffer (10 mM Tris-HCl pH 8.0, 1 mM EDTA, 500 mM NaCl, 1% Triton X-100, 0.1% SDS, 0.1% DOC, 1x Protease inhibitor (halt),1 mM PMSF), rotating at 4°C for 10 min during each wash. Carefully remove and discard supernatant each time. Wash the beads once with 1mL LiCl ChIP buffer rotating at 4°C for 10 min. Carefully remove and discard the supernatant. Wash the beads twice with 1mL TE, rotating at 4°C for 10 min. Carefully remove and discard the supernatant. Resuspend the beads in 150uL TE buffer. Add 10 uL 10% SDS, 5ul 5M NaCl, and Incubate for 6 hours or overnight at 65°C. Then Add 4.5 uL Proteinase K (20mg/ml) and 5 uL RNase A (.5 mg/ml). Mix well and incubate at 37 °C for 1-2 hours and then add 2 uL proteinase-K stop solution; mix it well and let it at room temperature for 5-10 mins. Extract samples with 150uL phenol-chloroform-isoamyl alcohol (25:24:1), vortex for 15 seconds, and spin for 10 min at room temperature, max speed. Take 120uL aqueous phase and back-extract the organic phase with 150uL TEN140 buffer (10mM tris-HCl pH 8.0, 1mM EDTA, 140 mM NaCl). Vortex for 15 seconds and spin for 10 min at room temperature at max speed. Take out 150uL of the aqueous phase and combine it with the aqueous phase from the above step (270uL aqueous phase total). Extract samples with 300uL chloroform, vortex for 10 seconds, and spin for 10 min at room temperature, max speed. The upper aqueous phase was pipetted to a new tube. Add 30uL 3M NaAcetate of pH 5.0 and 2uL glycogen (20mg/mL). Precipitate DNA with 900uL EtOH at -80°C for at least 1 hour. Spin for 25 min at 4°C, max speed. Carefully remove supernatant. Wash the pellet in 600uL 70% EtOH and spin the pellet for 10 min at 4°C at max speed. Carefully remove the supernatant and allow the pellet to air dry. Dissolve in 20uL 1X TE buffer and quantify the IP by Qubit and bioanalyzer before proceeding to library preparation.

### Library preparation

For library preparation, we have used NEBNext Ultra II DNA Library Prep Kits (E7645, New England Biolab NEB). We adopted the NEB protocol with some modifications. ChIP libraries were prepared from immunoprecipitated DNA for each sample. ChIP-ed fragmented DNA was diluted with 10mM Tris-HCl and adjusted the volume to 50ul in a 200ul PCR tube on a cold rack. Next, 3ul of End Prep Enzyme Mix and 7ul of End Prep Reaction buffer was added to the fragmented DNA and mixed well. Tubes containing the reaction mixture were placed in a PCR machine with the following setting: 20°C for 30 min, 65°C for 30 min, and 4°C holds. In the adapter ligation step, to the End, Prep Reaction Mixture, first added the 2.5ul of NebNext Adapter for Illumina mixed it well then added the 30ul of Ligation Master Mix, and 1ul of Ligation enhancer was incubated in a thermocycler at 20°C for 15 min. In the next step, USER enzyme (3ul) was supplemented to the reaction, and further, the mixture was incubated at 37°C for 15 min in a thermocycler with the heated lid set to > 47°C. At the size selection step, because of the low amount of initial material (2-5 ng), as recommended by the manufacturer, a size-selection step was not performed, and fragments were cleaned up to eliminate unligated adaptors by adding 87ul of Ampure XP beads (Beckman Coulter) and mixing it with the reaction volume. The mixture was incubated for 5 min at room temperature, and then separate the beads were using the magnetic stand, and the supernatant containing the short fragments was discarded. The beads were then washed twice with 200ul of freshly prepared 80%ethanol, air-dried for 5 min, and resuspended in 17ul of 10 mM Tris-Cl pH 8. Mix it well and incubate at room temperature for 2 minutes. Further, beads were separated using a magnetic stand, and transfer the supernatant containing DNA fragments was to a new 200ul PCR tube. For PCR amplification of each library, a PCR reaction mixture was prepared by adding 25ul of NEBNext Ultra II Q5 Master Mix, 5 ul of Universal PCR primer (i5 primer), and 5ul of Indexed primer (i7 primer) to the 15ul of the ChIP-Seq library. The PCR cycles were adjusted based on library concentration. The PCR reactions were run at 98°C for the 30s (98°C for 10 s, 65°C for 75 s) repeated 10 times, 65°C for 5 min, and 4°C holds. The PCR reaction was cleaned and size-selected using Ampure XP beads as follows: PCR reaction was resuspended in 45ul of AmpurXP beads, incubated at room temperature for 5 minutes, placed on the magnetic rack, discarded the supernatant containing small fragments, and kept the beads washed them twice with 200ul of freshly prepared 80%ethanol, remove the ethanol by placing on a magnetic rack. Air-dried them for 5 min and try to remove the traces of ethanol. Resuspended the beads in 33ul of 0.1x TE buffer and mix thoroughly and incubate them at room temperature for at least 2 minutes, then transfer the PCR tube to a magnetic rack. Then, transfer the supernatant containing the final library to a fresh PCR tube. We checked the size distribution on an Agilent bioanalyzer using a High Sensitivity DNA chip. And store the library at -20°C before sequencing the libraries.

### Hi-C library preparation

For Hi-C data, samples were collected the same as for ChIP-seq data. Initial processes of sample treatment are a little varied in Hi-C sample processing. After homogenization, the cells were resuspended in 1X PBS and then cross-linked with 1 % final formaldehyde concentration for 10 minutes at room temperature with end-over-end mixing. The crosslinking reaction was terminated with a quenching solution (200mM glycine) for 20 minutes at room temperature again with end-over-end mixing. The quenched cells were spun down and rinsed once with 1X PBS buffer. The cross-linked cells were homogenized and subsequently lysed in lysis buffer (10 mM Tris-HCl (pH 8.0), 10 mM NaCl, 0.2% NP40, and complete protease inhibitors and incubated for 20 minutes with end-over-end mixing. A high-speed spin was performed, then the supernatant was removed, and the pellet containing the nuclei was re-suspended with 150 μl 0.1% SDS and incubated at 65°C for 10 min. SDS molecules were quenched by adding 120 μl water and 30 μl 10% Triton X-100 and incubated at 37 °C for 15 min. To digest the chromatin, 150U of MboI and 30 μl 10x NEB buffer 2.1(50 mM NaCl, 10 mM Tris-HCl, 10 mM MgCl2, 100 μg/ml BSA, pH 7.9) were added. Then the reaction was incubated at 37 °C overnight. After the digestion, the MboI enzyme was inactivated at 65 °C for 20 min. Next, to biotinylate the free chromatin ends, 1 μl of 10 mM dTTP, 1μl of 10 mM dATP, 1 μl of 10 mM dGTP, 2μl of 5mM biotin-14-dCTP, 14 μl of water, and 4μl (40 U) Klenow were added to the reaction and incubated at 37 °C for 2 hours. In the next step, 663 μl water,120 μl 10x blunt-end ligation buffer (300 mM Tris-HCl, 100 mM MgCl2, 100 mM DTT, 1 mM ATP, pH 7.8), 100μl 10% Triton X-100 and 20 U T4 DNA ligase were supplemented to start proximity ligation. The proximity ligation reaction was incubated at 16 °C for 4 hours with end-over-end mixing. After four hours of ligation, the chromatin was reverse cross-linked by adding 200 µg/mL proteinase K (Thermo) at 65°C overnight. The next day, the free DNA was purified through QIAamp DNA Mini Kit (Qiagen) according to the manufacturer’s instructions. The purified DNA fragments were sheared in the range of ∼400 bp. The ligated and biotinylated Hi-C junctions were pulled down by using Dynabeads MyOne streptavidin according to manufacturers’ instructions. The beads were washed and re-suspended in elution buffer and used for library preparation using NEBNext® Ultra™ II DNA Library Prep Kit for Illumina (NEB). The amplified final libraries were sequenced on the Illumina HiSeq X Ten platform (San Diego, CA, United States) with 150PE mode.

### ChIP-seq data analysis

ChIP libraries of eleven histone modification marks, including input controls in the embryonic stage, sexed larvae, and adult tissues in *Dpse*, were sequenced. 150PE ChIP-seq reads were mapped to the *Dpse* de novo chromosomal level assembly using bwa-mem with minimal overlap of 30 bp (q<=20). Sorting and indexing of bam files were performed by using SAMtools^60^. Illumina adapters were trimmed using trimmomatic^61^. Peaks were called with MACS2^62^ on each replicate and merged using cat, bedtools ^63^ sorts (Version_v2.29.2) followed by BEDtools merge with 10 bp option (-f BAMPE -g dm -n out -q 0.01) were used. Coverage files were generated with deepTools (Version_3.5.1)^64^ bamCompare, using binsize of 10 bp (--bs 10 --*minMappingQuality 10 --normalizeTo1x {EFFECTIVE_GENOME_SIZE}*”). Duplicated reads and reads with a mapping quality < 10 were removed, the 1x-normalization method was used, and the X chromosome was ignored for normalization. IP over Input log2-fold ratios were calculated for all hPTMs using bamCompare with settings “*-bs 10 –scaleFactorsMethod SES*’’. Heatmaps were generated using deepTools adopted commands computeMatrix and plotHeatmap with default parameters. The parameters like bamCompare and bamCoverage from deepTools were used to create normalized coverage of bigWig files, and genome-browser snapshots were generated using IGV^65^. For motif analysis, we used MEME^66^ (Version_5.2.0) adopted parameters: -nmotifs 3 -dna -revcomp -mod zoops -objfun classic -markov_order 0 -p 4. We used chromHMM (Ernst and Kellis 2012) to call the chromatin states genome-wide using a 15-state chromatin model with the following steps; we prepared and gave the annotation files such as genes, TSS, TES, intron, exons, and TEs, and then ChIP-seq hPTMs along with input control with single cell type option. First, we proceeded with the binarization step and called the regions of significant enrichment genome-wide (per chromosome) at 200bp resolution. Next, combine the enrichment profile of each chromosome from the previous step and learn models with different numbers of chromatin states. Post-learning models were compared to check the consistency between the models and previously published models. 15-state chromatin models were more consistent and reliable learned from our set of hPTMs. However, our model achieved good segregation of each chromatin state and compatibility with the annotation pattern.

### Total RNA-seq analysis

Total RNA alignment was performed using RSEM ^67^ against the reference transcript sequences and gene annotation using bowtie2 with default parameters. Reads were counted per transcript and summed for each gene using the “rsem-calculate-expression” function in the RSEM package. Tissue-specific log2-fold change gene expression levels were calculated using the DESeq2 package ^68^. To test whether there are differences in gene expression levels during development, a TAU score was calculated (a higher TAU value means more stage/tissue specificity). Differentially expressed genes are listed in Supplementary Data.

### Hi-C data bioinformatics analysis

*Dmel* and *Dpse* sexed Hi-C data sets were produced for embryonic stage 12, 3^rd^ instar male larvae, head, and testes using MboI as a restriction enzyme carried out as described by^11^. Paired-end fastq reads were mapped, and Hi-C matrices were generated and normalized using HiC-Pro and HiC-explorer ^69,70^. Forward and reverse reads were trimmed and mapped separately to the UCI_Dpse_MV25 reference genome considering only chromosomes 2, 3, and X. We excluded chrY and chrDot from the analysis for the reason that either they are mainly heterochromatic or very short in size. Thus, they may not be comparable at chromosomal level analysis, like interaction frequency decay with distance. We used bowtie2 (version 2.3.5.1) to run Hic-Pro alignments with default parameters except for q-value cutoff, which was set to 10 and --very-sensitive, where -L 30 and --score-min L,-0.6,-0.2 was used. We first used Hi-C pro utilities to Identify the genomic locations of each DpnII restriction site, the following command was used (digest_genome.py -r DpnII). Non-digested, self-ligated fragments were removed and reads were initially counted for each valid DpnII fragment to generate a fragment-level Hi-C object. Restriction fragment level Hi-C objects were then merged into bins of equal size at different resolutions, including 5, 10, 25, and 100 kb resolutions. Finally, the binned matrices were normalized using the iterative correction method ^71^. Moreover, to remove non-informative read pairs, we excluded the first two diagonals during normalization (interactions at distances 0 or 1 bin). For ICE normalization of Hi-C contact maps, we used the iterative_mapping module^71^ (iced version 0.5.2) in HiC-Pro for aligning reads to the reference genome, making the assumption of equal visibility of each fragment.The following filtering parameters were adopted in ICE pipeline: --filter_low_counts_perc 0.02 and --filter_high_counts_perc 0, we used --max_iter 100, --eps 0.1 and --remove-all-zeros-loci, --output-bias 1, --verbose 1. We also filtered for duplicated or non-informative read pairs associated with PCR artifacts that were discarded during normalization. Only valid pairs involving two different restriction fragments were used to build the ICE–normalized contact matrices. The biological replicates from sex-sorted Hi-C embryos were mapped separately and merged into their corresponding samples. To further analyze these ICE-normalized matrices, we convert them to h5 format using ‘hicConvertFormat’ from HiCExplorer^70^. Hi-C matrix visualization was performed using ‘hicPlotMatrix‘ or ‘pyGenomeTracks‘ for the specified regions. TADs were called using ‘hicCallTADs’ and the following parameters were adopted ‘--correctForMultipleTesting fdr --numberOfProcessors 30 --minBoundaryDistance 5000 --thresholdComparisons 0.01 −delta 0.05 --step 5000’. That way, we obtained the TAD boundary positions and genome-wide TAD insulation scores. Conserved and non-conserved TABs between different developmental stages (embryo and larvae) and adult tissues (head and testes) were calculated (by extending the TABs by 2 kb) using bedtools intersect (version v2.29.2), and the minimum overlap cutoff was set to 50% (-f 0.5). Tissue or stage-specific TABs are defined as only differentially appearing in one tissue or stage but disappearing from others. To check detectable differences in the Hi-C interaction profiles between tissue. We compared the log2-fold ratio of contact frequencies of normalized Hi-C matrices binned at 25 kb resolution shown for a representative 5Mb region.We compared the Hi-C signal ratio in mid (500kb) and long-range (3MB) distances. We performed fold-change comparisons between male vs. female contact interaction frequencies of each bin were then calculated using FAN-C^72^. commands were used: fanc compare -c fold-change -l -Z –I –tmp to create fold-change matrices and then plot using the command: fancplot X:5mb-10mb -p triangular -m 2.5mb -c RdBu_r -vmin -5 -vmax 5. Heatmap showing the pairwise difference (deltas) of the rate of Hi-C signal decay (the slope coefficient). The slope coefficients were computed considering distances <400 kb (≤5.6 in log_10_ distance). Hicexplorer^70^ and FANC^72^ were used as HiC data visualization tools and for other downstream analyses like TAD calling. The first eigenvector (PC1) corresponding to active (A) and inactive (B) compartments was computed using FANC after the removal of heterochromatic chromosome ends. Corrected Hi-C matrices of 25 kb resolution were used to call compartments. To switch the orientation of PC1 values, where positive values correspond to the active compartment (A) and negative values correspond to the inactive compartment (B), we used GC contents. In the end, we verified the PC1 orientation for each chromosome to overlay with active and inactive histone modification mark ChIP-seq data. The saddle plot shows preferential interactions of active vs. active (AA) in red and inactive vs. inactive (BB) regions in blue color. The bar plot on top shows the cutoffs used for binning regions by the corresponding EV entry magnitude. We used a common approach to summarize the average pairwise Hi-C contacts between a set of selected genomic loci. hicAggregateContacts from HicExplorer with corrected Hi-C matrices and settings “*--numberOfBins 60 --vMin 1 --vMax 2 --range 300000:5000000 --plotType 3d –avgType mean –chromosomes –transform obs/exp* ” to plot aggregated pairwise Hi-C contacts between TEs and between TEs and H3K27acpeaks.The window of size n is projected both on the row and column as indices, i and j, respectively. In our case, the binning resolution of the Hi-C matrix was 5 kb and n = 60. Therefore, the examined window was of size ±15 or 30 kb surrounding any given pair of loci.

